# Resistance to heavy metals and genome sequencing of ESBL-producing and non-ESBL-producing Escherichia coli

**DOI:** 10.1101/2025.05.28.656662

**Authors:** Ma Wenyi, Yuan Tianshuai, Duan yu, Xu yue, Sui saihui, Sun Han, Yin YiFei, Ji Hua

## Abstract

The present study aimed to examine the impact of heavy metals on the resistance levels exhibited by ESBL-producing and non-ESBL-producing *Escherichia coli*. The minimum inhibitory concentration (MIC) of six heavy metals on two types of E. coli was determined using the broth microdilution method. Heavy metal resistance genes were detected via PCR amplification, and the gene structures were analyzed through whole genome sequencing. The findings indicated that ESBL-producing E. coli strains exhibited significantly higher levels of resistance to Cu, Zn, and Cd compared to non-ESBL-producing strains. Additionally, ESBL-producing strains harbored various plasmid-mediated heavy metal resistance genes. The results of WGS demonstrated that ESBL-producing strains carried plasmids encoding 148 heavy metal resistance genes, and the resistance to certain antibiotics (e.g., chloramphenicol) in E. coli exhibited a significantly negative correlation with resistance to specific heavy metals, including Mn and Cd (*p* < 0.05). The present study hypothesizes the presence of synergistic mechanisms between heavy metal resistance and antibiotic resistance in ESBL-producing E. coli, including the dual functionality of the efflux pumping system, the synergistic regulation of the pressure response system, and the physical barrier effect of biofilm formation. The experimental findings revealed the link between heavy metals and antibiotic resistance, highlighting the importance of environmental monitoring and control of heavy metal pollution. Further studies are warranted to explore the mechanisms by which heavy metals promote the transfer of genes associated with antibiotic resistance in order to formulate effective intervention strategies to combat this issue.

## Introduction

*Escherichia coli* (E. coli)(1) is a key pathogenic microorganism responsible for the development of mastitis in dairy cows, a disease that significantly affects the dairy industry and the economic output of dairy farming. The dissemination of extended-spectrum β-lactamase (ESBL)-producing E. coli has been recognized as a significant global public health concern(2). At present, the primary strategy for controlling mastitis is antibiotic therapy, in which β-lactam antibiotics are extensively employed to manage mastitis and other bacterial diseases(3). However, these antibiotics have been demonstrated to exert greater selective pressure, contributing to the occurrence and dissemination of ARGs (antimicrobial resistance genes). The present study demonstrated that ESBL genes are principally associated with a broad spectrum of splicing-type plasmids(4). Notably, the propagation of antibiotic resistance may not only be driven by the antibiotic itself but also by non-antibiotic stressors in the environment (e.g., heavy metals), which may similarly exacerbate resistance through synergistic effects(5). In the context of microbial communities subject to selective pressure from heavy metals, there is a high probability of co-selection for heavy metal resistance genes. Furthermore, the presence of heavy metals (e.g., Cu^2+^, Zn^2+^) in livestock farming environments raises concern, given that these metals may contribute to the evolution of multi-drug resistance in conjunction with antibiotic exposure through various mechanisms, including an increased frequency of horizontal transfer of antibiotic resistance genes (ARGs)(6), alterations to the structure of the microbial community(7), and the induction of ARGs(8). The potential origins of these synergistic effects may be rooted in animal husbandry practices.

Heavy metals and antibiotics have been observed to co-exist across various environmental matrices, including the gastrointestinal tract, animal feces, and poultry farms. Yan et al(9) established that the effects of different heavy metals on bacteria markedly vary, with swine-sourced isolates of ESBL-producing *Escherichia coli* exhibiting high levels of resistance to Cu, Zn, Cd, and Cr. For instance, the introduction of zinc oxide and copper sulfate to pig feed has been demonstrated to enhance post-weaning performance(10). However, it is worthwhile emphasizing that the European prohibits the administration of high levels of zinc oxide as of June 2022 (EMA 2017). Furthermore, research has indicated that the addition of heavy metals, such as Cu^2^ ^+^ and Zn^2^ ^+^, to animal feed at low concentrations, balances trace mineral levels, increases antimicrobial effects, and promotes healthy growth. In a study examining soil samples from Scottish soils, Knapp et al.(11) identified a positive correlation between numerous ARGs and soil copper levels. Additionally, they noted significant correlations (*p* < 0.05) between specific ARGs and chromium, nickel, lead and iron. Heavy metals are characterized by their persistence and capacity for accumulation within diverse components of the ecosystem. Sahin et al.(2) documented that ESBL-producing E. coli isolated from chicken meat exhibited high levels of resistance to heavy metals. Furthermore, the presence of genes associated with heavy metal resistance was detected within E. coli. Furthermore, Chen et al.(12) pointed out that the bacterium LSJC7 demonstrated a high level of resistance to copper and zinc. In addition, the study established a correlation between heavy metal concentration and bacterial resistance to antibiotics. Noteworthily, the application of poultry manure to the soil as a fertilizer is an effective method for recycling nutrients. However, the accumulation of these trace minerals in environments such as animal manure or soil may lead to residual contamination, possibly ascribed to the production and spread of AMR through triggering co-option or co-regulation of resistance genes(13).

The co-primary objectives of this study were to detect genotypes and phenotypes of heavy metal resistance in β-lactamase-producing and non-β-lactamase-producing strains and to compare the results of the former with those of the latter. In order to establish the relationship between heavy metal resistance genes (HMRGs) and antibiotic resistance genes (ARGs), it is necessary to undertake a comprehensive analysis of the whole genome sequence of two multidrug-resistant *Escherichia coli* strains. This approach will facilitate a more detailed understanding of the structure of resistance genes and an in-depth analysis of the characteristics of heavy metal resistance in these two distinct E. coli strains. It is hypothesised that this information may contribute to the monitoring and control of the spread of this bacterial population.

## Materials and methods

### 2.1 Materials

Brain Heart Infusion broth (BHI) medium, Eosin-methylene blue medium, Brain Heart Infusion agar (BHI) medium, Müller-Hinton agar were purchased from Qingdao HaiBo Biotechnology Co. 6 × Sampling buffer, dNTPs, 10 × buffer, Tap DNA polymerase, DNA markers, and Gold View Nucleic Acid dye were purchased from Tiangen Biochemical Technology Co. Tiangen Biochemical Technology Co.

### 2.2 Strain screening

A total of 766 samples were collected from five dairy farms in Shihezi, Urumqi, and Yili from August 2020 to November 2021 by the laboratory. Among these farms, one was of a small-scale nature, and four operated on a commercial scale. Furthermore, significant variations in herd sizes were noted across the farms, ranging from approximately 100 animals on smaller farms to over 500 animals on larger commercial farms. The sampling strategy encompassed 570 raw milk samples and 196 environmental samples, including 79 cow manure, 22 sewage, 11 fencing, 13 water troughs, 27 soil, 23 cow carcass, 13 feed, and 8 hand swab samples. E. coli was isolated using two distinct agar media, namely eosin-methylene blue agar (Difco TM) and MacConkey agar (Difco TM). Typical E. coli colonies were biochemically confirmed via the indole test. Individual E. coli colonies were collected from each positive sample, placed in 20% glycerol, and stored at 80 °C in a refrigerator. A total of 133 ESBL-producing E. coli strains and 31 non-ESBL-producing E. coli strains were selected to investigate their resistance to heavy metals. All strains are included in the collection of Food of the Institute of Microbiology at the University of Shihezi (China).Bacteria were grown in brain-heart infusion broth (BHI) (Qingdao HaiBo Biotechnology, China) or on BHI agar (BHI with 1.5% agar) at 37 °C for 24 or 36 h, depending on the strain.

### 2.3 Heavy metal minimum inhibitory concentration (MIC)

The MIC of metal ions against ESBL-producing E. coli was determined using the agar microdilution method described by the Clinical and Laboratory Standards Institute(14). The six heavy metals utilized in this study comprised Cu (CuCl₂· 2H₂O), Cr (CrCl₃·6H₂O), Co (CoCl₂), Cd (CdCl₂), Zn (ZnSO₄), and Mn (MnCl_2_). Isolates demonstrating resistance to a minimum of three heavy metals were designated as multi-resistant strains. Each experiment was conducted in triplicate. ATCC29522 served as the reference standard strain.

### 2.4 Amplification and detection of heavy metal resistance genes

E. coli DNA was extracted and subsequently utilized as the template for Polymerase Chain Reaction (PCR) amplification. The PCR reaction system (25 µL) comprised 12.5 µL of Taq Master Mix (2×), 0.5 µL of each specific primer (20 µM), 10.5 µL of ddH_2_O, and 1 µL of the bacterial template. These components were mixed homogeneously prior to amplification. The PCR products were subjected to analysis on a 1.2% (w/v) agarose gel operating at a voltage of 120 V for 30 minutes. The amplification results were then visualized using a gel imager. Positive controls derived from ESBL-producing E. coli isolates were selected for amplification. The positive controls were confirmed by sequencing of the amplicons by (Qingdao Sangong Biotechnology Co., Ltd.). To ensure reproducibility, all experiments were independently performed at least twice. The primer sequences and PCR conditions are listed in Table 1.

**Table 1.**
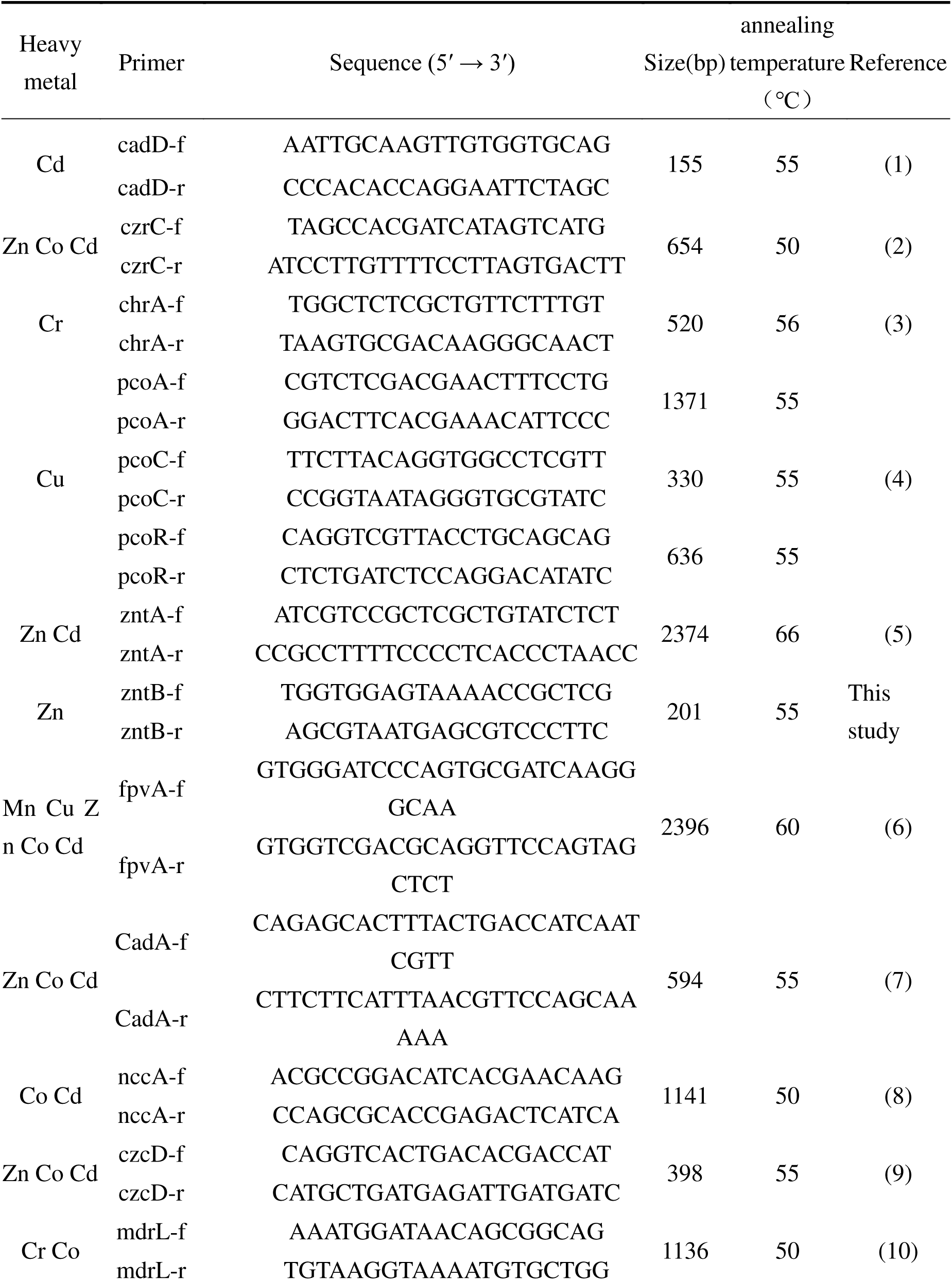
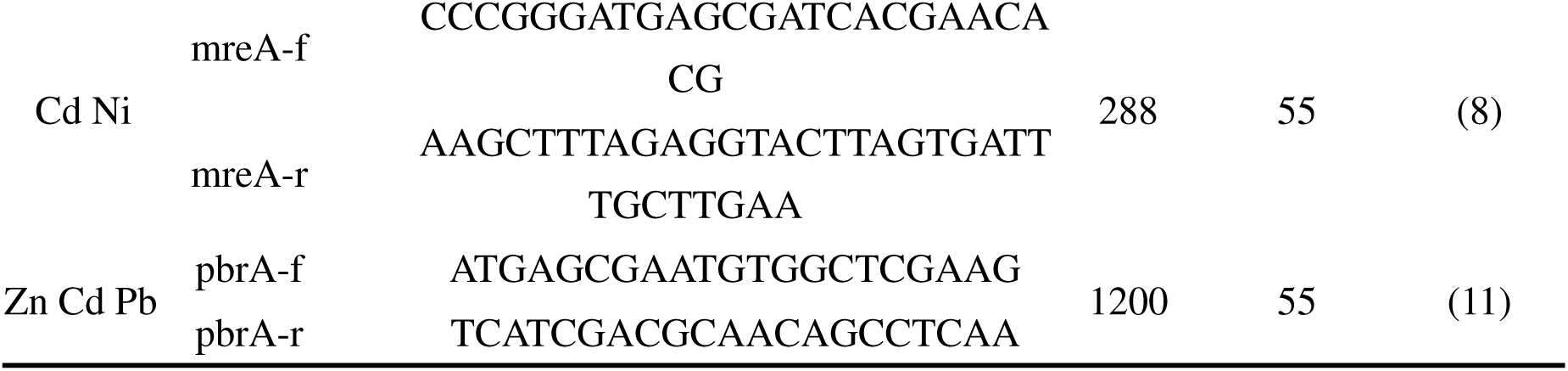
The following primers have been utilised in this study for the purpose of synthesising DNA from some of the metal resistance genes under investigation:

**Table 2.** The following investigation focuses on the comparison of heavy metal resistance (MIC) in ESBL-producing E. coli with that observed in non-ESBL-producing E. coli.

| Heavy metals | Type | MIC( μ L/L) |  |  |  |  |  |  | MIC<sub>50</sub> | MIC<sub>90</sub> | Resistance % | P.value |
| --- | --- | --- | --- | --- | --- | --- | --- | --- | --- | --- | --- | --- |
|  |  | 256 | 512 | 1024 | 2048 | 4096 | 8162 | > 8162 |  |  |  |  |
|  |  | Number of strains |  |  |  |  |  |  |  |  |  |  |
| Cu | ESBLs | 4 | 14 | **106** | 9 | - | - | - | 1024 | 2048 | 85.80% | <0.01 |
|  | non-ESBLs | - | 16 | **14** | - | - | - | - | 1024 | 1025 | 46.70% |  |
| Zn | ESBLs | - | 1 | 38 | **66** | 27 | - | - | 2048 | 4096 | 69.40% | <0.01 |
|  | non-ESBLs | - | - | 25 | **5** | - | - | - | 2048 | 2048 | 16.70% |  |
| Mn | ESBLs | - | - | - | 15 | 77 | **40** | - | 8162 | 8162 | 30.50% | <0.01 |
|  | non-ESBLs | - | - | - | 7 | 8 | **16** | - | 8162 | 8162 | 53.30% |  |
| Cd | ESBLs | 7 | 1 | - | 7 | 14 | **103** | - | 8162 | 8162 | 76.80% | <0.01 |
|  | non-ESBLs | - | - | - | - | 15 | **16** | - | 8162 | 8162 | 53.30% |  |
| Cr | ESBLs | 2 | 11 | 58 | **51** | 10 | - | - | 2048 | 4096 | 45.50% | <0.01 |
|  | non-ESBLs | 2 | 11 | 58 | **3** | 10 | - | - | 2048 | 4096 | 43.30% |  |
| Co | ESBLs | 5 | 58 | **64** | 4 | - | - | - | 1024 | 2048 | 50.70% | <0.01 |
|  | non-ESBLs | 1 | 18 | **12** | - | - | - | - | 1024 | 1024 | 40% |  |

### 2.5 Correlation between antibiotic resistance and heavy metal resistance

The relationship between metal resistance and antibiotic resistance in ESBL-producing E. coli was investigated using the chi-squared (x^2^) test of independence or Fisher’s exact test. Statistical analyses were performed using SPSS v.21 software. A p-value of less than 0.05 was considered statistically significant.

### 2.6 Comparative whole genome sequencing of heavy metal-resistant strains

The present study investigated the phenotypes and genotypes of the experimental strains for heavy metal resistance, with one ESBL-producing E. coli strain and one non-ESBL-producing E. coli strain selected for whole genome sequencing.

#### 2.6.1 Gene assembly

Total DNA was initially extracted using the CTAB method, a widely employed extraction kit. Subsequently, the DNA was fragmented into fragments ranging from 300 to 500 base pairs (bp) using a Bioruptor (Diagenode, Belgium). Sequencing libraries were generated using the VAHTS Universal DNA Library Preparation Kit for Illumina (Vazyme, ND 627) in accordance with the manufacturer’s instructions. This process encompassed the utilization of an articulator ligation, magnetic bead purification, PCR amplification, and cyclization. Finally, the libraries were sequenced on a Novaseq-6000 instrument, employing a PE read length of 150 base pairs. Trimmomatic (version 0.36) was used to filter articulators, and low-quality reads from the β-lactamase mechanism.

#### 2.6.2 Gene Annotation & Analysis

To obtain comprehensive gene function information, gene function annotations were performed in major databases, including UniProt, GO(15), Pfam,COG(16), KEGG(17), and KEGG Pathway, and the top 20 most annotated structural domains were graphically displayed. A statistical summary of the genes annotated for each structural domain is reported based on the annotations of the Pfam structural domains. Potential plasmids were identified in the sequencing data as contiguous clonal lines with high copy numbers, indicative of their extraction as separate entities. Thereafter, a search was conducted for homologs of plasmid maintenance genes, followed by comparisons with known plasmids. Genomic islands were identified using the Island Viewer 4 software, whilst potential AMR genes were identified by conducting a BLAST search. The BacMet metal resistance gene database (version 2.0), a repository of experimentally confirmed sequences, was utilized to identify potential metal resistance genes through the implementation of a BLAST search algorithm. Exocytosis pump genes were identified using two methods. Firstly, the Bac Met database was utilized as a comprehensive reference source. (“BacMet: antibacterial biocide and metal resistance genes database | Nucleic Acids Research | Oxford Academic,” n.d.)Secondly, genome annotations were manually examined. Lastly, metal resistance and antibiotic resistance genes were referenced with IslandViewer 4(18) results to assess the presence of resistance gene clusters in putative genomic islands.

The WGS data of all 2 *Escherichia coli* strains were submitted to the National Center for Biotechnology Information (NCBI), with BioProject accession numbers SUB15259191 ID: PRJNA1250 and SAMN47966503 ID: PRJNA1250975, respectively.

## Results

### 3.1 MIC of heavy metals on ESBL-producing E. coli isolates

The present study revealed the differential inhibitory effects and resistance characteristics of six heavy metals (Cu, Zn, Mn, Cd, Cr and Co) by analyzing the minimum inhibitory concentration (MIC) distributions and the resistance rates among ESBL-producing E. coli isolates.

Specifically, Cu demonstrated significant sensitivity (*p* < 0.05) at moderate MIC levels (1024 μg/L, 106 strains), with only a limited number of strains requiring higher concentrations (2048 μg/L, 9 strains) for inhibition. The MIC distributions of Zn and Cd were skewed towards high concentration ranges (Zn: 2048-4096 μg/L, 93 strains; Cd:8162 μg/L, 103 strains) Mn demonstrated elevated MIC values (4096-8162 μg/L, 117 plants), indicative of its dual functionality as both a toxic substance and an essential micronutrient. In contrast, the MIC values for Cr and Co were within the median range of the spectrum (1024-2048 μg/L for Cr). Likewise, the MICs for Cr and Co were within the intermediate range (Cr: 1024-2048 μg/L, 109 strains; Co: 512-1024 μg/L, 122 strains), suggesting the presence of heterogeneity in the tolerance mechanisms of microorganisms to these elements. Interestingly, ESBL-producing strains demonstrated significantly higher levels of resistance to Cu, Zn, and Cd compared to non-ESBL-producing strains (*p* < 0.05), indicating that multi-resistance mechanisms may enhance heavy metal tolerance through gene co-transfer (e.g., plasmid-mediated metal resistance genes). Moreover, the resistance exhibited by ESBL-producing E. coli to Cu, Zn, and Cd was significantly higher compared to non-ESBL-producing strains (*p* < 0.05), signaling the occurrence of a synergistic evolution between the multi-drug resistance and heavy metal tolerance.

ESBL-producing E. coli from bovine milk exhibited a high degree of resistance to heavy metals, with the strongest resistance observed against Cu and Cd (resistance ratios of 50/50 and 48/50 for both, with an average resistance of 100% and 96.7%, respectively), followed by Zn (72.4%) and Co (47.2%). Strains of fecal origin demonstrated a comparatively high level of resistance to Cu (52.6%) and Zn (73.7%) and exhibited a substantially lower level of resistance to Mn (19.1%) and Cd (35.9%) compared to alternative sources (*p* < 0.05). The analysis of environmental samples revealed complete resistance to Cd (25/25, 100%) and very low resistance to Mn (3.8%). In addition, non-ESBL-producing strains derived from bovine milk demonstrated minimal resistance, with higher resistance exclusively noted for Mn and Cd at levels of 52.1% and 51.6%, respectively. A comprehensive analysis of heavy metals unveiled that copper exhibited the highest level of resistance (85.8%), followed by cadmium (76.8%), whereas manganese demonstrated the lowest level of resistance (30.5%). Furthermore, the resistance exhibited by homologous strains to various heavy metals exhibited significant variation. For instance, ESBL-producing strains demonstrated 55.6% and 51.5% higher resistance to Cu and Cd, respectively, compared to Cr.

### 3.2 Heavy metal resistance genes

Among ESBL-producing E. coli, the highest rates of resistance to Cu and Cd were observed in strains derived from bovine milk (Cu: 50/50 = 100%, Cd: 60/62 ≈ 96.8%), whilst strains of environmental origin were 100% resistant to Cd (25/25), and fecal origin displayed a significantly lower resistance (Cu: 22/42 ≈ 52.4%, Cd: 15/42 ≈ 35.7%) (*p* < 0.05). Genetic testing uncovered that ESBL-producing strains harbored multiple plasmid-mediated heavy metal resistance genes, such as czrC (36/50, 72%), pcoR (41/50, 82%), and pbrA (44/50, 88%) in cow milk, and czrC (19/25, 76%) and pbrA (21/25, 84%) were prominently detected in environmental sources. Conversely, non-ESBL-producing strains exhibited a predominance of chromosomally encoded genes, such as fpvA. At the same time, broad-spectrum resistance genes (e.g., fpvA) were detected in 83.9% (26/31) of cow milk samples, whereas plasmid-encoded genes (e.g., czrC, zntB) were absent. Furthermore, CadA (7/50) and mreA (15/50) were exclusively detected in cow milk-derived ESBL-producing strains, czcD (3/25) in environmental samples, and mdrL (17/42) in fecal strains, indicating source-specific distributions. Overall, the prevalence of ESBL-producing E. coli resistance genes was higher than in non-ESBL-producing E. coli, both in terms of species and abundance. Lastly, the distribution of genes originating from cow milk and environmental sources from R14 was predominantly characterized by exogenous heavy metal contamination, while fecal sources exhibited a greater reflection of selective pressures exerted by the host’s internal environment.

In contrast, non-ESBL-producing E. coli exhibited chromosome-dependent broad-spectrum resistance genes (e.g., fpvA) in cow milk, lacked plasmid-mediated multiple resistance, and displayed more conserved resistance mechanisms. A cross-comparison of the two sources indicated a high prevalence of resistance genes in ESBL-producing strains derived from cow milk and environmental samples, highlighting the accelerating role of anthropogenic activities (e.g., farming and industry) on the evolution of bacterial resistance.

### 3.3 Correlation between antibiotic resistance and heavy metal resistance

In combination with earlier antibiotic resistance experiments on E. coli(19), resistance to chloramphenicol in E. coli exhibited a significantly negative correlation with heavy metal resistance, specifically Mn and Cd (*p* < 0.05). Furthermore, resistance to Cd was significantly correlated with multidrug resistance (*p* < 0.05). However, this phenomenon was not universally observed, indicating that strains exhibiting elevated antibiotic minimal inhibitory concentrations (MICs) do not invariably correspond to elevated MICs for heavy metals. Conversely, this finding suggests the presence of diversity and variability in the mechanisms underlying antibiotic and biocide resistance. Strains exhibiting elevated MIC values for specific heavy metals may not be categorized as clinically resistant due to the absence of breakpoints that delineate resistance to heavy metals. Consequently, a definitive association between antibiotic resistance and heavy metal resistance could not be established. Thus, further investigation was undertaken to explore genomic associations underlying these resistance phenotypes.

### 3.4 GC view Chart

Utilizing the third-generation sequencing platform of Nanopore, ESBL-producing E. coli R14 yielded 89,519 filtered and validated reads, with an average length of 10,171.58 base pairs. On the other hand, the non-ESBL-producing E. coli R150 generated 46,561 filtered, validated reads, with an average length of 9,924.96 bp. These results corroborated that the sequencing output is suitable for subsequent bioinformatics analysis of high-quality sequencing data. Complete genome sequences were obtained by splicing the genome sequences of the two strains, revealing circular chromosomes with lengths of 3,431,884 bp and 4,440,328 bp, and GC contents of 43.66% and 53.93% for R14 and R150, respectively. Furthermore, the genome contained 4,221 and 5,009 genes, accounting for approximately 79.5% and 81.47% of the whole genome. The distribution of protein lengths is illustrated in Figure 4, with the majority of proteins exhibiting genomic lengths of at least 1,000 amino acids, accounting for over 30% of the total. The circular genome profiles of R14 and R150 are presented in Figure 4ab, offering valuable insights into their genomic organization.

### 3.5 Summary of gene annotations

R14 (ESBL-producing strains): the total number of annotated genes was relatively low, which may be indicative of the reduced complexity of the genome or the incomplete annotation of some genes. The annotation numbers of COG and KEGG were 3833 and 4240, respectively, suggesting specific capabilities in their functional classifications and metabolic pathways. R150 (non-ESBL-producing strains): the total number of annotated genes was relatively high (e.g., NR: 4566, Swiss-Prot: 4350), indicating a more comprehensive functional annotation of its genome. An analysis of core functional databases (COG, GO, KEGG) revealed the presence of 3,878, 3,753 and 3,395 annotated genes, respectively, indicating the maintenance of metabolic and functional pathways.

This gene annotation map is a multidimensional framework for elucidating gene function, constructed by integrating annotation information from six authoritative databases: NR, Swiss-Prot, Pfam, COG, GO, and KEGG. The sequence coverage provided by NR is complemented by the accuracy of Swiss-Prot and GO, while KEGG, Pfam, and COG provide further information related to pathways, structural domains, and evolutionary relationships. These databases collectively form the ‘pyramid’ of functional annotation. In the context of antibiotic resistance, KEGG annotations in R14 (4,240) exhibited higher compared to those in R150 (3,395), suggesting the potential enrichment of resistance genes (e.g., β-lactamase-related genes) within its metabolic pathway. The divergent patterns observed in the annotations of the GO database (R14: 4,127 vs. R150: 3,753) might be associated with molecular functions (e.g., hydrolase activity) or biological processes (e.g., enhanced antibiotic catabolism). Besides, Pfam analysis of protein families and structural domains revealed that R14 exhibited a higher number of annotations (4458) compared to R150 (4244), implying that the ESBL-producing strain R14 may possess a greater number of unique structural domains. Evidently, conserved domains associated with heavy metal resistance genes were identified.

Regarding R14 (ESBL-producing strains), the total number of annotated genes was comparatively low, indicative of the reduced complexity of the genome or the incomplete annotation of some genes. The number of annotations in the COG and KEGG databases was 3833 and 4240, respectively, suggesting the presence of specific adjustments in functional classification and metabolic pathways. Conversely, the total number of annotated genes for R150 (non-ESBL-producing strains) was comparatively higher (e.g., NR: 4566, Swiss-Prot: 4350), indicating a more comprehensive functional annotation of its genome. The number of annotated genes in the core functional databases (COG, GO, KEGG) were 3878, 3753, and 3395, respectively, indicating the conservation of metabolic and functional pathways.

The Kyoto Encyclopedia of Genes and Genomes (KEGG) is a comprehensive database that integrates information on biological genomes, metabolic pathways, and compounds. This resource enables researchers to systematically analyze biological processes, thereby providing a robust framework for exploring the intricate mechanisms underlying cellular activities(17).

The two strains exhibited analogous core genetic mechanisms linked to DNA replication and transcription yet diverged in specific functional pathways. For instance, strain R14 exhibited elevated activity in horizontal gene transfer-related pathways, including transposon and recombination. This finding indicates that R14 is highly likely to acquire new genes by carrying ESBL-encoding plasmids. In response to external stimuli, R14 primarily relies on metabolic pathways such as membrane transport (ko02010) and the two-component system (ko02020), which have been demonstrated to play a pivotal role in bacterial responses to external stresses and in performing antibiotic efflux. Conversely, the function of strain R150 appears to be oriented towards pathways associated with nutrient uptake and environmental adaptation.

Concerning metabolic characterization, R150 showed a stronger correlation with pathways such as cell division and carbohydrate metabolism, indicating its active growth state and enhanced efficiency in energy utilization. Conversely, R14 was predicted to allocate a greater proportion of its resources to survival mechanisms in response to stress, such as the oxidative phosphorylation pathway (ko00190). Regarding antibiotic resistance, the ESBL phenotype of R14 was observed to be closely related to the high expression of β-lactamase-related genes and efflux pumps. The necessary metabolic support for these functions is provided by pathways such as the ABC transporter protein (ko02010) and penicillin/cephalosporin biosynthesis (ko00311) pathways.

Nevertheless, it is imperative to acknowledge the presence of a discernible metabolic trade-off strategy within R150. The rationale behind this is as follows:In a similar manner, a total of 3,833 protein sequences of ESBL-producing E. coli R14 were annotated in the COG database, accounting for 85.94% of the genome. Similarly, the non-ESBL-producing E. coli R15 had a total of 3,878 protein sequences annotated, representing 84.91% of the genome (Fig. 7). The annotated units of R14 were largely focused on carbohydrate transport and metabolism (421), amino acid transport and metabolism (368), transcription (325), and energy production and conversion (295), with similar major categories observed in R150. A greater number of protein sequences were annotated in the DNA repair unit R14 (155) compared to R150 (146). Finally, 421 genes were annotated in R14 and 378 genes in R150 in the energy production and conversion unit.

### 3.6 Removable element gene island sets (GIs) and plasmids

The sequence length of the R14-exclusive plasmid pR17.4942_71k was 87,315 base pairs (bp) with a guanine-cytosine (GC) content of 53.79% (Figure 4). The backbone exhibited(GenBank submition: CP100720.1) the characteristic features of the IncF type(20), which has been documented to regulate plasmid replication, facilitate horizontal transfer, and regulate plasmid repression and stabilization. Characterization of Inc/rep plasmid types in E. coli strains revealed that IncFIB-type plasmids were prevalent in both ultra-broad-spectrum β-lactamase (ESBL) and non-ESBL-producing strains. However, in both strains, the presence of the gene was exclusively detected in the ESBL-producing R14 strain, whereas in non-ESBL-producing isolates, it was identified as IncFIC(FII). Of note, strains harboring the blaCTX-M-15 gene and resistant to quinolones, aminoglycosides, and sulfonamides were of the IncF plasmid type. As indicated by the BacMet database annotation, 148 HMRGs were identified on the plasmid unique to R14 in the present study, including HMRG isoforms for polymetallic resistance, such as copper, arsenic, nickel, iron and other common metals. Furthermore, exocytosis pump genes and removable elements were annotated on the plasmid, enhancing the potential for the transfer of antibiotic resistance.

A search was conducted for genomic islands in the strains, with a particular emphasis on the presence of antibiotic resistance genes (ARGs) and metal resistance genes located on these islands as potential indicators of co-resistance. R14 and R150 contained 20 and 16 genomic islands, respectively. These islands were identified in the ESBL-producing E.coli R14 strain. More importantly, R14 possessed a 21,977 bp genomic island (GI02) unique to this strain. This island contains genes that confer resistance to metals, including nickel and cobalt, as well as proteins involved in the export of nickel and cobalt ions and genes that encode proteins that facilitate membrane transport. This island is depicted in Figure 2.

**Fig. 1.**
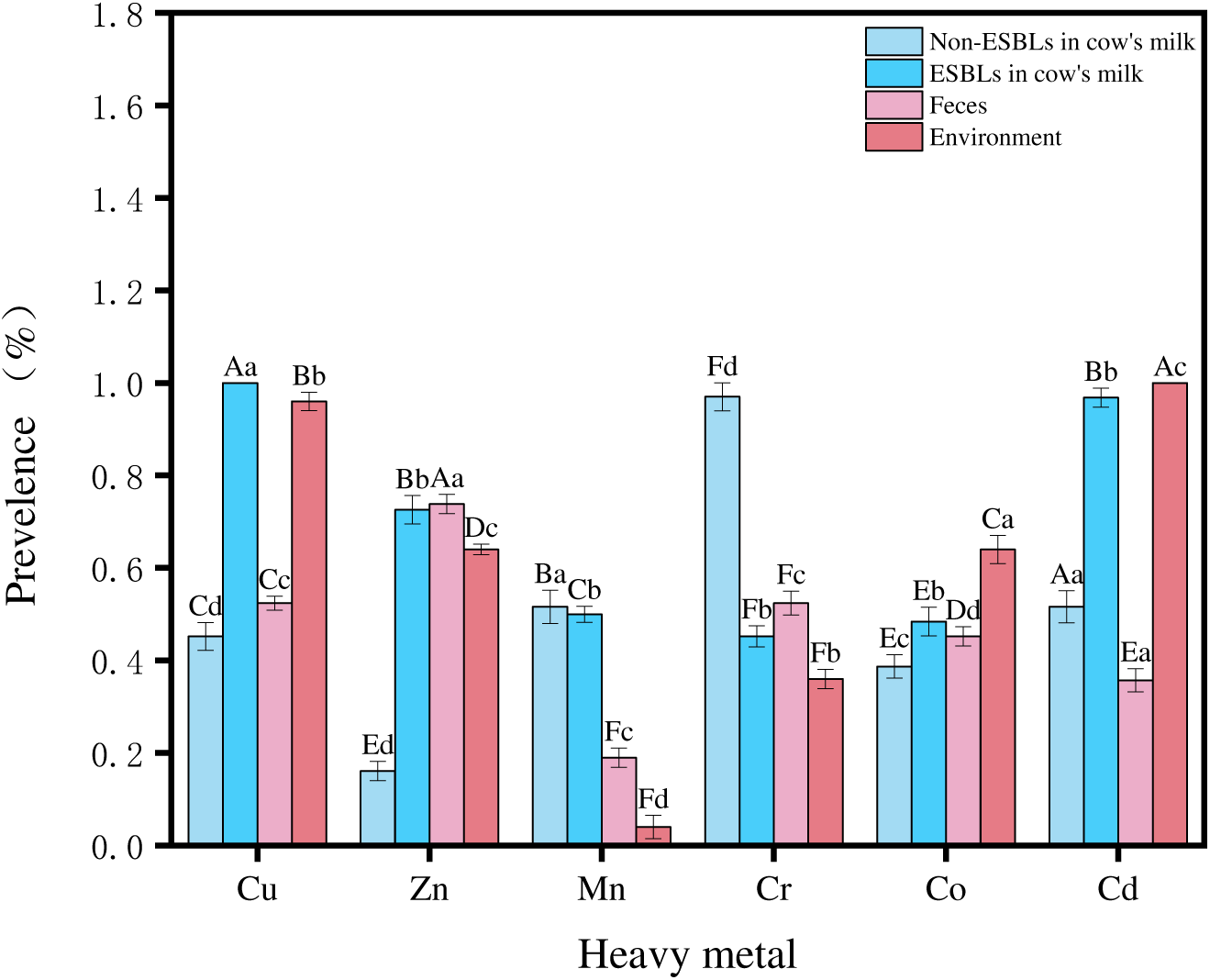
The present study investigates the prevalence of heavy metal resistance in E. coli from various sources. The investigation focuses on the prevalence of heavy metal resistance to copper(Cu), zinc(Zn), manganese(Mn), cadmium(Cd), chromium(Cr) and cobalt(Co) in ESBL-producing E. coli from milk, non-ESBL-producing E. coli, faeces and E. coli isolated from the bovine farm environment. The utilisation of different letters denotes statistically significant differences between groups within groups (*P* < 0.05).

**Fig. 2.**
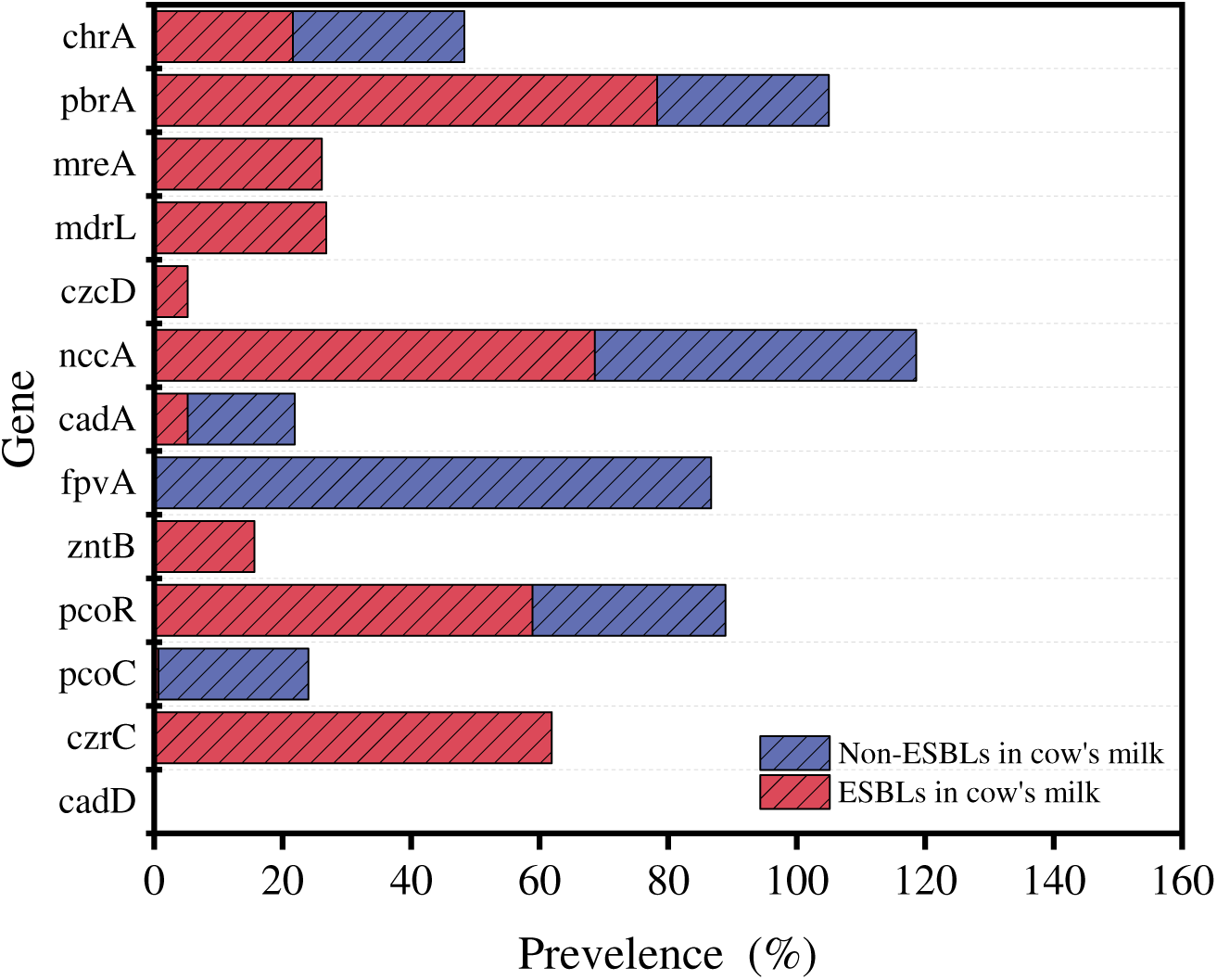
The results from a heavy metal resistance-mediated PCR screen.

**Fig. 3.**
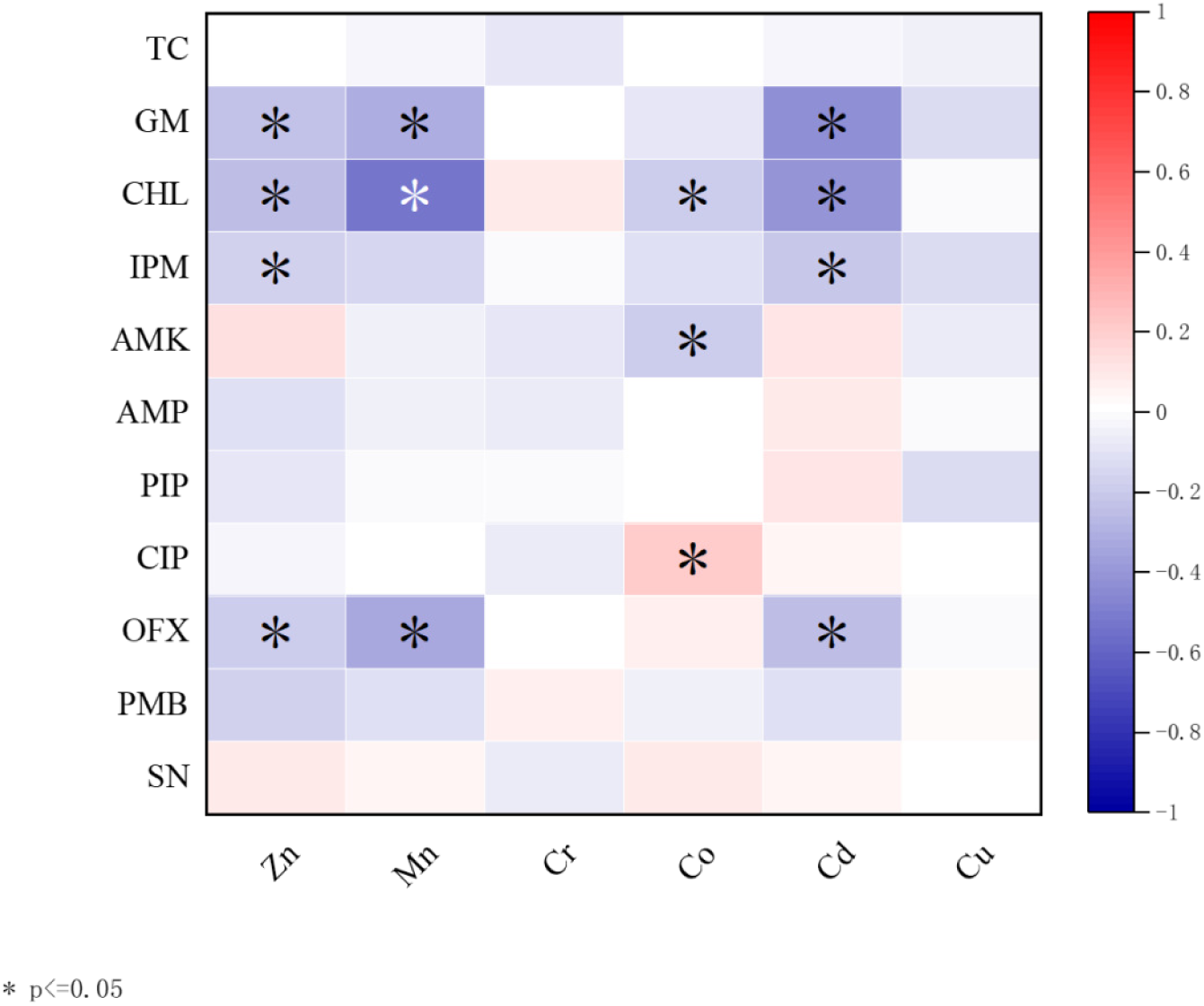
Correlation between antibiotic resistance and heavy metal resistance The abbreviations in the diagram are TC ÿTetracyclineĀÿGM ÿGentamicinĀÿCHL ÿChloramphenicolĀÿIPM ÿImipenemĀÿAMK ÿAmikacinĀÿAMP ÿAmpicillinĀÿ PIP ÿPiperacillinĀÿCIP ÿCiprofloxacinĀÿOFX ÿOfloxacinĀÿPMB ÿPolymyxin BĀÿSN ÿSulfonamideĀ; The utilisation of different letters denotes statistically significant differences between groups within groups (*P* < 0.05).

**Fig. 4.**
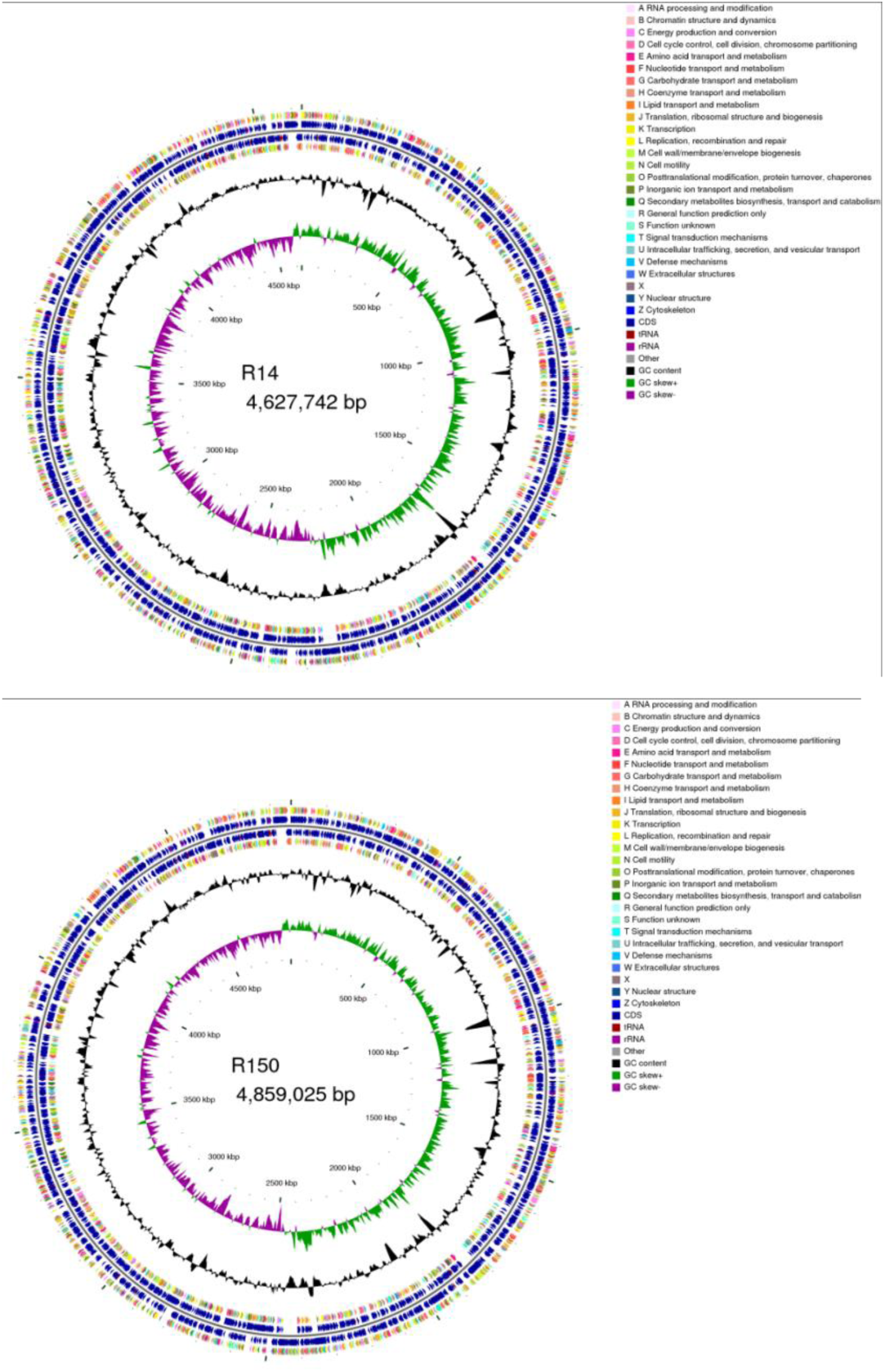
GCview chart The first and fourth circles are CDS on the positive and negative strands, and different colours indicate different COG functional classifications; the second and third circles are CDS, tRNA and rRNA on the positive and negative strands, respectively; the fifth circle is the GC content, and the outward part of the circle indicates that the GC content of the region is higher than the average GC content of the whole genome, and the higher the peak indicates the larger difference with the average GC content; the sixth circle is the GC-Skew value, and the inward part indicates that the GC content of the region is lower than the average GC content of the whole genome; the specific algorithm is GC-Skew value. The higher the peak value, the greater the difference with the average GC content; the sixth circle is GC-Skew value, the specific algorithm is G-C/G+C, which can assist in determining the leading and lagging strand, generally the leading strand GC skew>0, the lagging strand GC skew<0, and it can also assist in determining the replication start point (cumulative offset) and end point (cumulative offset). Minimum) and end point (maximum cumulative skew), especially important for circular genomes; the innermost circle is the genome size marker.The first graph is the gene circle map for R14 and the second is the gene circle map for R150.

**Fig. 5.**
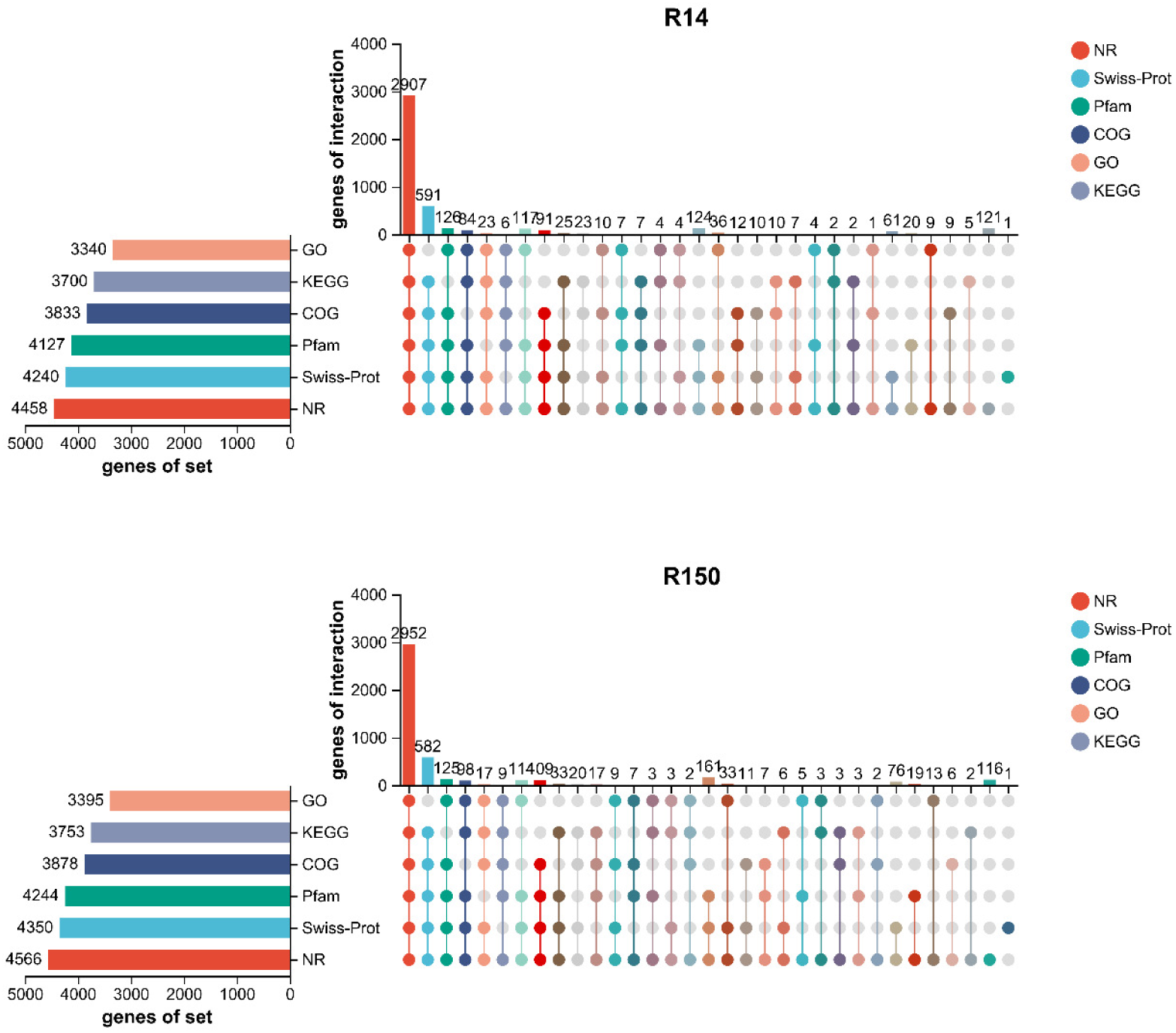
Statistical annotation of the Upset plot The horizontal bar chart on the left represents the total number of genes annotated in each database. In the central matrix, a single dot indicates the unique genes annotated by a specific database corresponding to the left bar, while connected dots represent genes commonly annotated by multiple databases. The vertical bar charts respectively display the counts of corresponding unique/shared genes.

**Fig. 6.**
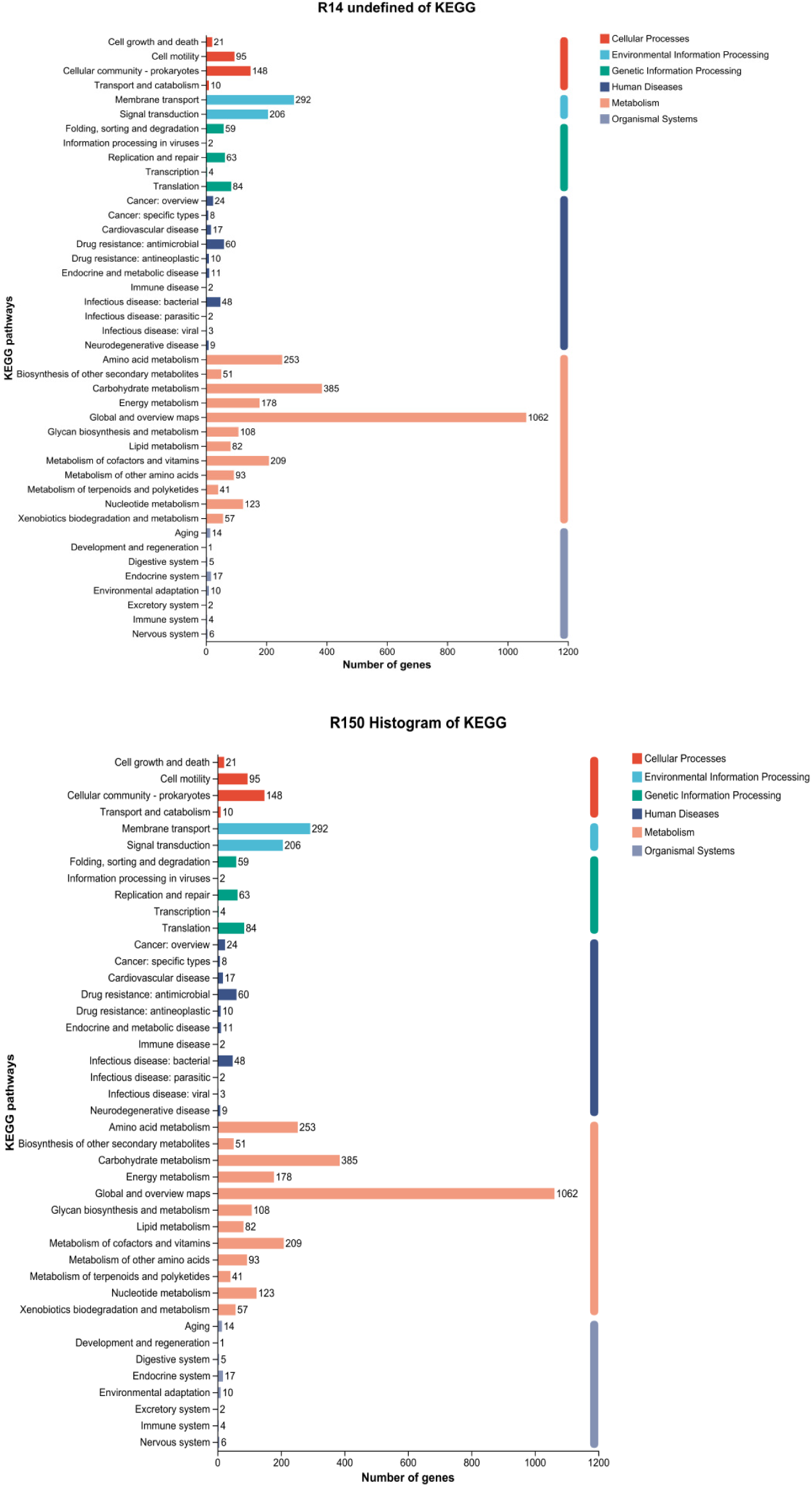
The KEGG bar chart uses the vertical axis to represent the level 2 hierarchical classification of KEGG pathways, while the horizontal axis indicates the number of genes annotated under each classification. The colors of the bars represent the level 1 hierarchical classification of KEGG pathways. The rightmost bar displays the gene counts under different level 1 classifications. Since the same gene may be annotated to multiple level 2 classifications, deduplication is performed when calculating the gene counts for level 1 classifications.

**Fig. 7.**
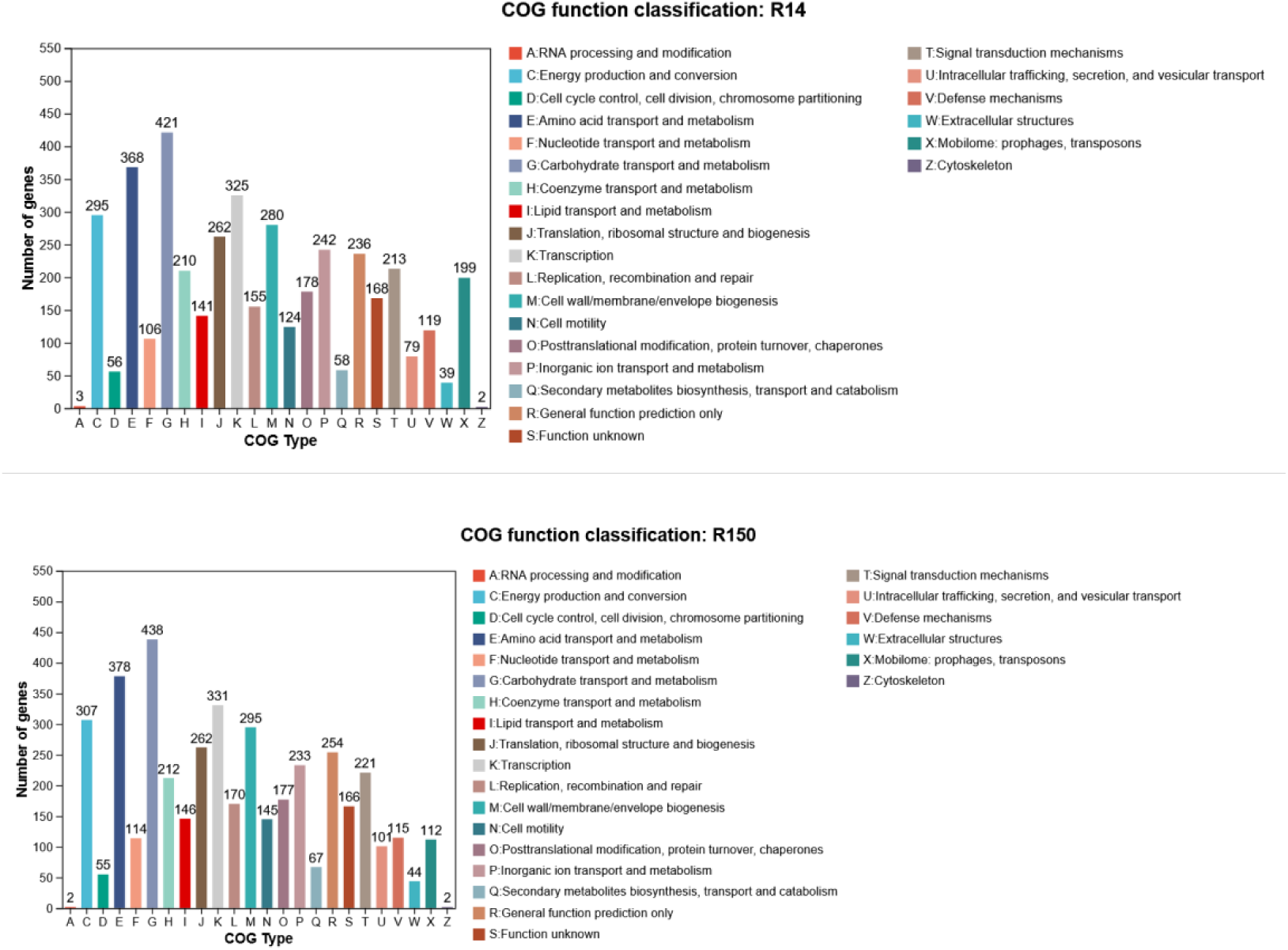
COG Classification Statistics Bar Chart The horizontal axis represents different COG categories, and the vertical axis represents the number of genes. For functional descriptions of each specific COG category, please refer to the legend on the right.

**Fig. 8.**
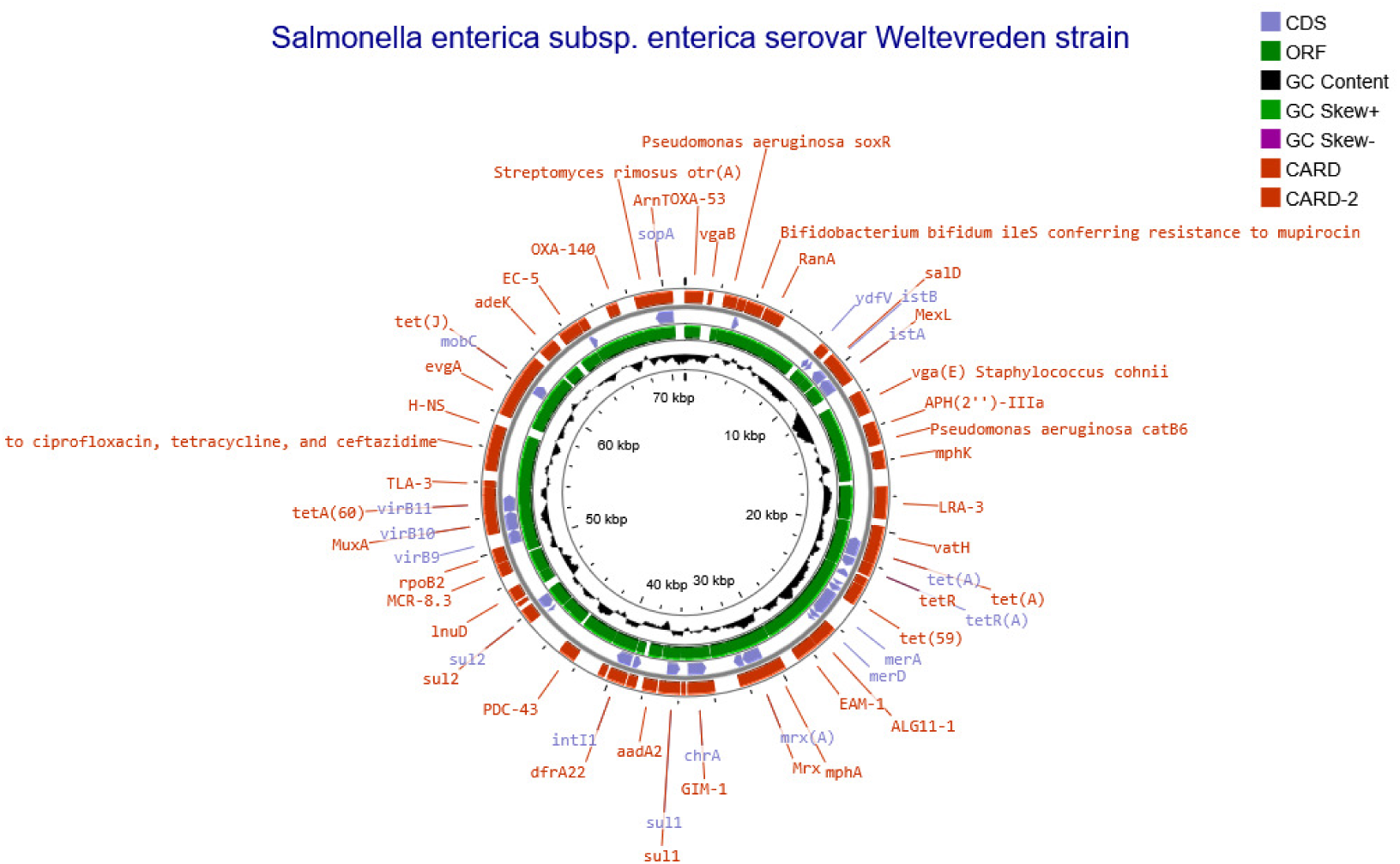
Plasmid genetic circle diagram Red represents antibiotic resistance gene annotation; green represents mobile genetic elements (MGEs); blue represents coding sequences, the regions of the genome capable of encoding proteins; black represents GC content.

**Fig. 9.**
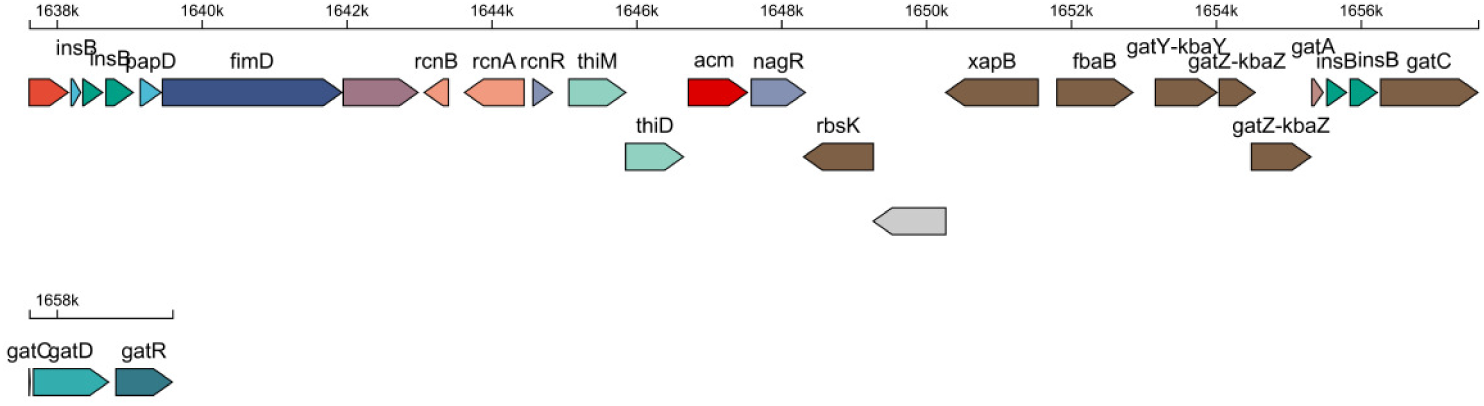
Linear Genetic Island Map GI02 Genomic island linear map, where each arrow represents a gene, the length of the arrow indicates the gene length, the direction denotes whether the gene is encoded by the sense strand or antisense strand, the color represents its COG functional classification, and uncolored arrows signify genes without annotated COG functions.

## Discussion

This study demonstrated that ultra broad-spectrum β-lactamase (ESBL)-producing E. coli exhibited significant resistance (p<0.05) to heavy metal environments. The distribution of resistance in descending order was as follows: Cu (85.8%) > Cd (76.8%) > Zn (69.4%) > Co (50.7%) > Cr (45.5%) > Mn (30.5%). This finding suggests that Cu and Cd exert the most significant selective pressure on microorganisms among heavy metals (Table 1). At the same time, the findings of the present study demonstrated that the minimum inhibitory concentration (MIC) for Cu and Zn were consistent with those reported by Deus et al. (2017)(21). The distribution of MIC values for all heavy metals was narrowly concentrated within one to two consecutive concentration intervals. Nevertheless, regarding genotypes, Yang et al.(20) discovered that ESBL-producing E. coli isolated from German chicken farms possessed the highest number of resistance genes, encompassing zntA, which conferred resistance to zinc and cadmium. This phenomenon may be attributed to the substantial addition of zinc to poultry feed. In the case of highly resistant metals such as Cu, Cd, Zn, etc., E. coli has been hypothesized to excrete heavy metal ions from cells through an active exocytosis mechanism or use an intracellular chelation process to bind heavy metal ions to specific substances, thus limiting their toxicity.

Conroy et al. (22) identified an RND-type multiprotein system in E. coli, termed the metal-exporting system CusCFBA, responsible for detoxification during metal stress. In the context of low- to medium-resistance metals, such as Co, Cr, Mn, and others, bacteria may rely on redox balance regulation to maintain the stability of the intracellular environment. Alternatively, they may achieve tolerance to heavy metals by ingesting nutrients in a manner that mimics metal uptake. However, the evolution of resistance pathways may be constrained by numerous factors(23). Importantly, a substantial difference in MIC distribution was observed between Cu (concentration-sensitive) and Cd (threshold-dependent), indicating differences in the mechanisms of tolerance even for metals exhibiting comparable levels of resistance.

Conversely, the observed differences in heavy metal resistance between ESBL-producing and non-ESBL-producing E. coli may be attributable to: i) the synergistic effect of the efflux pump. ESBL-producing strains have been observed to enhance the active efflux of heavy metals such as Cu, Zn and Cd, potentially through the carriage of both metal resistance genes (e.g., czc, mer) and antibiotic resistance genes (e.g., _bla_ESBL_) on plasmids. Secondly, the differential expression of metal-binding proteins may also account for these variations(23). The high resistance to copper may be related to the induction of metallothionein expression (24)and the higher efficiency of antioxidant systems (e.g., SOD enzymes) in ESBL-producing strains. These systems can mitigate the cytotoxicity of metal ions(25).

Furthermore, heavy metal resistance in ESBL-producing E. coli from diverse sources exhibited variability, with environmental samples demonstrating complete resistance to Cd (100%), potentially attributable to ongoing selection pressure resulting from industrial pollution. In contrast, the low resistance to Mn (3.8%) may be indicative of its reduced environmental concentration or toxicity threshold. The high resistance of fecal strains to Zn (73.7%) may be related to the widespread use of Zn additives in animal feed(26). It is worthwhile pointing out that Cr and Co exhibited less than 50% resistance across all sources, which may be related to their lower bioavailability. Within-group comparisons revealed heterogeneity in resistance to different heavy metals among strains from the same source. For instance, resistance to Cd was 96.2% higher than Mn in environmental samples, suggesting adaptive differentiation of strains to specific heavy metals. Heavy metal resistance is generally associated with a multifaceted interplay among strain origin, metal exposure history, and genetic background.

Moreover, a significant correlation was identified between heavy metal resistance and antibiotic resistance, along with their associated ARGs (*p* < 0.05), in line with the findings of Deng et al(27)., which investigated the co-occurrence of heavy metal and antibiotic resistance in Salmonella isolated from retail food of animal origin and noted a significant correlation (*p* < 0.05). According to Puangseree et al(28).ESBL-producing *Escherichia coli* isolated from pork swine farms also demonstrated significant resistance to disinfectants, heavy metals, and antibiotic resistance. The experimental data presented herein conjointly suggest that ESBL-producing E. coli exhibit enhanced heavy metal resistance, a phenomenon that may be attributable to the co-selection mechanism of antibiotic resistance genes and heavy metal resistance genes. Earlier studies have established(11) a positive correlation between the abundance of antibiotic-resistance genes in the environment and elevated concentrations of antibiotics and heavy metals. Similarly, numerous studies have concluded that the co-localization of heavy metal resistance genes with antibiotic resistance genes is prevalent in clinical isolates of ESBL-producing *Escherichia coli*(29) (8). This phenomenon is most likely a consequence of evolution driven by co-selection pressures arising from combined exposure to heavy metals and antibiotics in agricultural or medical settings(30). The complexities of this issue are further compounded by the distinct expression levels of heavy metal and antibiotic resistance genes in ESBL-producing E. coli. The efflux pump system exerts dual effects. R14 has been found to be significantly enriched in pathways related to environmental information processing, and the ‘membrane transport’ (e.g., ABC transporter protein, ko02010) pathway has been shown to exhibit enhanced efflux pump activity (31). ABC transporter proteins have been evinced to be effective in the extracellular excretion of β-lactam antibiotics, as well as in the excretion of heavy metal ions, including Zn²⁺ and Cu²⁺. Meanwhile, the up-regulation of the expression of efflux pump genes (e.g., acrAB-tolC) can confer bacterial tolerance to both antibiotics and heavy metals, thereby offering a survival advantage in complex environments(32). Secondly, the role of synergistic regulation by stress-responsive systems must be considered. R14 has been demonstrated to be active in the ‘two-component system’ (ko02020) and ‘oxidative phosphorylation’ (ko00190) pathways, which reflects its strong stress-responsive and energy metabolism abilities. The two-component system (e.g., PhoPQ, BaeSR) has been demonstrated to govern the expression of genes associated with antibiotic and heavy metal resistance(33). Furthermore, it has been observed to exhibit a high degree of flexibility in adjusting the expression levels of related genes in response to variations in environmental stress. Oxidative phosphorylation plays a decisive role in supplying ATP, which is essential for energy-consuming processes such as efflux pumps(27). This, in turn, provides sufficient energy substrate to maintain the multiple resistance phenotype. Furthermore, the presence of certain heavy metals (e.g., Cu⁺) has been observed to induce oxidative stress through the generation of reactive oxygen species (ROS), thereby activating antioxidant pathways. Prior investigations have postulated that the SOD enzymes of R14 may be more efficient in scavenging ROS, which may indirectly enhance its survivability under heavy metal stress environments. Functional annotations based on the COG database also revealed that R14 was significantly enriched in genes related to DNA repair and may be more adept at repairing heavy metal-induced DNA damage (e.g., free radical-induced strand breaks) and enhancing genomic stability. Thirdly, the role of biofilm formation cannot be overlooked. ESBL-producing E. coli have been observed to be more closely associated with genes related to biofilm formation(34). As a specialized structure, biofilms can form a physical barrier on the cell surface, which effectively limits the penetration of heavy metal ions and attenuates intracellular toxicity. In addition, Tripathi et al(31). discovered that efflux pumps significantly promote biofilm formation and maturation through various mechanisms, including the efflux of extracellular polymeric substances and molecules involved in population quenching, to assist in establishing the biofilm matrix. ESBL-producing strains are typically characterized by enhanced biofilm-forming ability, a process that can be facilitated(35) by population sensing systems (e.g., luxS) or genes implicated in extracellular polysaccharide synthesis (e.g., wca clusters), which eventually increase their resistance to heavy metals.

Herein, strain R14 was significantly enriched in horizontal gene transfer-related pathways (e.g., transposons, recombination, and other pathways) involved in genetic information processing. This observation suggests the possibility of multiple resistance plasmids in strain R14. ESBL genes such as blaCTX-M and blaTEM are frequently located on removable genetic elements, such as plasmids or integrons. These plasmids also harbor heavy metal resistance genes, including copper, cadmium, and arsenic resistance genes, such as copA and czcABC(36). The concurrent dissemination of ARGs and MRGs via MGEs in response to co-selection pressure exerted by antibiotics and heavy metals remains a significant public health concern. The co-transfer phenomenon has been demonstrated in several ecological niches, including the human gut. Rajput et al(37). determined that exposure to Cu2+ concentrations exceeding 5 μg/L increased plasmid-mediated splicing transfer of ARGs within bacterial genera among *Escherichia coli* strains.

The IncFIB plasmid harbored by strain R14 exhibits a unique and complex biological profile. This plasmid is notable for its possession of typical antibiotic resistance genes, including blaCTX-M-15, in addition to a substantial enrichment of metal resistance genes. The integration of resistance modules, including transposons and integrons, culminates in the aggregation of multiple resistance genes and the formation of ‘ resistance islands’(38). Furthermore, in the presence of exogenous plasmids, IncFIB plasmids possess the capacity to elevate their resistance phenotype through a process referred to as ‘gene complementation’(39). A representative example is the AcrAB-TolC exocytosis pump system encoded on the plasmid, which is capable of not only participating in the exocytosis of antibiotics but also in effluxing heavy metal ions from cells, demonstrating its multifunctionality in conferring resistance. Furthermore, R14 possesses a unique genomic island, GI02, which encodes nickel/cobalt efflux proteins, as well as a regulatory transport system that largely enhances its adaptability to heavy metal environments. This synergistic resistance phenotype may be closely related to the long-term evolutionary strategy of the IncF plasmid. In the context of environmental selective pressure, such as exposure to heavy metals in medical wastewater, IncF plasmids can establish a sophisticated resistance advantage. This is achieved by strategically positioning metal resistance modules in close proximity to antibiotic resistance genes via transposons or integrons(40).

## Conclusion

This study demonstrated that the resistance of ultra-broad-spectrum β-lactamase (ESBL)-producing *Escherichia coli* to heavy metals, such as Cu, Zn, and Cd, was substantially greater than that of non-ESBL-producing strains. Moreover, the results revealed that the prevalence of resistance genes was chiefly characterized by plasmid-mediated efflux pumping systems, including pcoAB, czrAB, and other mobile genetic elements. In contrast, non-ESBL-producing strains predominantly relied on chromosomally encoded conserved genes. The co-selection of antibiotic and heavy metal resistance is facilitated by the dual function of efflux pumps, synergistic regulation of two-component systems, and plasmid co-localization. Besides, differences in resistance between strains of different origins were closely related to heavy metal exposure resulting from human activities. The present study corroborates the notion that sub-inhibitory concentrations of heavy metals can accelerate the transfer of drug-resistant genes, highlighting the necessity for enhancing the regulatory oversight of heavy metal contamination in the environment to impede the transmission of drug-resistant bacteria at the ‘animal-environment-human’ interface. Overall, this study offers a theoretical reference for the prevention and management of drug resistance.

## Author Contributions

Ma Wenyi: Conceptualisation, data curation, and writing-original draft. Yuan Tianshuai: Data curation, methodology, and validation. Duan yu: Data curation, formal analysis and methodology. Xu yue: Investigation, methodology, and validation. Sui saihui:Methodology, Investigation, Formal analysis, Data curation. Sun Han: Formal analysis, methodology. Yin YiFei:Formal analysis, Data curation. Ji Hua: Funding acquisition, supervision, and editing.

## Data Availability Statement

Data are contained within the article or supplementary material.

## Acknowledgements

This study was supported by grants from the National Natural Science Foundation of China (NSFC)-Regional Fund Project, Prevalence Characteristics of ESBLs-producing *Escherichia coli* in the Dairy Production Chain in Xinjiang Region and the Mechanism of Transmission of Typical Drug Resistance Genes (32260621); Corps Excellence Youth Project (Ji Hua); Basic Research Programme of Shihezi University (MSPY202403).

## References

1. Wileman BW, Thomson DU, Olson KC, Jaeger JR, Pacheco LA, Bolte J, Burkhardt DT, Emery DA, Straub D. 2011. Escherichia coli O157:H7 Shedding in Vaccinated Beef Calves Born to Cows Vaccinated Prepartum with Escherichia coli O157:H7 SRP Vaccine. Journal of Food Protection 74:1599–604.

2. Sahin S. 2022. Disinfectant and heavy metal resistance profiles in extended spectrum β-lactamase (ESBL) producing Escherichia coli isolates from chicken meat samples. Int J Food Microbiol.

3. Hille K, Felski M, Ruddat I, Woydt J, Schmid A, Friese A, Fischer J, Sharp H, Valentin L, Michael GB, Hörmansdorfer S, Messelhäußer U, Seibt U, Honscha W, Guerra B, Schwarz S, Rösler U, Käsbohrer A, Kreienbrock L. 2018. Association of farm-related factors with characteristics profiles of extended-spectrum β-lactamase- / plasmid-mediated AmpC β-lactamase-producing Escherichia coli isolates from German livestock farms. Veterinary Microbiology 223:93–99.

4. Vitt AR, Sørensen AN, Bojer MS, Bortolaia V, Sørensen MCH, Brøndsted L. 2024. Diverse bacteriophages for biocontrol of ESBL- and AmpC-β-lactamase-producing 产 ESBL 和 AmpC β-内酰胺酶的多种噬菌体的生物 防治 E. coliE.杆菌. iScience 27:108826.

5. Yu M-F, Chen L, Liu G, Liu W, Yang Y, Ma L. 2025. Metagenomic deciphers the mobility and bacterial hosts of antibiotic resistance genes under antibiotics and heavy metals co-selection pressures in constructed wetlands. Environmental Research 269:120921.

6. Baker-Austin C, Wright MS, Stepanauskas R, McArthur JV. 2006. Co-selection of antibiotic and metal resistance. Trends in Microbiology 14:176–182.

7. Zhang R, Gu J, Wang X, Li Y. 2020. Antibiotic resistance gene transfer during anaerobic digestion with added copper: Important roles of mobile genetic elements. Sci Total Environ 743:140759.

8. Sun H, Cai S, Liu H, Li X, Deng Y, Yang X, Cao S, Li W, Chen H. 2023. FgSdhC Paralog Confers Natural Resistance toward SDHI Fungicides in Fusarium graminearum. J Agric Food Chem 71:20643–20653.

9. Yan M, Wang W, Jin L, Deng G, Han X, Yu X, Tang J, Han X, Ma M, Ji L, Zhao K, Zou L. 2024. Emerging antibiotic and heavy metal resistance in spore-forming bacteria from pig manure, manure slurry and fertilized soil. Journal of Environmental Management 371:123270.

10. Zhang Y, Gu AZ, Cen T, Li X, He M, Li D, Chen J. 2018. Sub-inhibitory concentrations of heavy metals facilitate the horizontal transfer of plasmid-mediated antibiotic resistance genes in water environment. Environmental Pollution 237:74–82.

11. Knapp CW, McCluskey SM, Singh BK, Campbell CD, Hudson G, Graham DW. 2011. Antibiotic Resistance Gene Abundances Correlate with Metal and Geochemical Conditions in Archived Scottish Soils. PLoS ONE 6:e27300.

12. Chen S, Li X, Sun G, Zhang Y, Su J, Ye J. 2015. Heavy Metal Induced Antibiotic Resistance in Bacterium LSJC7. Int J Mol Sci 16:23390–23404.

13. Puangseree J, Jeamsripong S, Prathan R, Pungpian C, Chuanchuen R. 2021. Resistance to widely-used disinfectants and heavy metals and cross resistance to antibiotics in *Escherichia coli* isolated from pigs, pork and pig carcass. Food Control 124:107892.

14. Singer AC, Shaw H, Rhodes V, Hart A. 2016. Review of Antimicrobial Resistance in the Environment and Its Relevance to Environmental Regulators. Front Microbiol 7.

15. Iram D, Sansi MS, Puniya AK, Meena S, Vij S. Draft genome sequence of methicillin-resistant Staphylococcus aureus strain D1418m22 isolated from human wound pus. Microbiol Resour Announc 12:e00409–23.

16. Liu Q, Wang H, Zhu W, Peng S, Zou H, Zhang P, Li Z, Zhang Z, Fu L, Qian Z. 2024. Determination of extracellular proteinase in *L. helveticus Lh191404* based on whole genome sequencing and proteomics analysis. International Journal of Biological Macromolecules 276:133958.

17. Kanehisa M. 2004. The KEGG resource for deciphering the genome. Nucleic Acids Research 32:277D – 280.

18. Bertelli C, Laird MR, Williams KP, Lau BY, Hoad G, Winsor GL, Brinkman FSL. 2017. IslandViewer 4: expanded prediction of genomic islands for larger-scale datasets. Nucleic Acids Res 45:W30–W35.

19. Huang S, Tian P, Kou X, An N, Wu Y, Dong J, Cai H, Li B, Xue Y, Liu Y, Ji H. 2022. The prevalence and characteristics of extended-spectrum β-lactamase Escherichia coli in raw milk and dairy farms in Northern Xinjiang, China. International Journal of Food Microbiology 381:109908.

20. Yang S. 2020. Presence of heavy metal resistance genes in Escherichia coli and Salmonella isolates and analysis of resistance gene structure in E. coli E308. Journal of Global Antimicrobial Resistance.

21. Deus D, Krischek C, Pfeifer Y, Sharifi AR, Fiegen U, Reich F, Klein G, Kehrenberg C. 2017. Comparative analysis of the susceptibility to biocides and heavy metals of extended-spectrum β-lactamase-producing *Escherichia coli* isolates of human and avian origin, Germany. Diagnostic Microbiology and Infectious Disease 88:88–92.

22. Conroy O, Kim E-H, McEvoy MM, Rensing C. 2010. Differing ability to transport nonmetal substrates by two RND-type metal exporters: Substrate specificity of metal RND exporters. FEMS Microbiology Letters no-no.

23. Zhang Y, Gu AZ, Cen T, Li X, He M, Li D, Chen J. 2018. Sub-inhibitory concentrations of heavy metals facilitate the horizontal transfer of plasmid-mediated antibiotic resistance genes in water environment. Environmental Pollution 237:74–82.

24. Maraschi AC, Adorno HA, Gonçalves YC, Souza IC, Monferrán MV, Wunderlin DA, Fernandes MN, Monteiro DA. 2024. Effects of metallic dust on Nile tilapia: Exploring the relationship between metal bioaccumulation, metallothionein levels, and oxidative stress responses. Science of The Total Environment 956:177423.

25. Lemire JA, Harrison JJ, Turner RJ. 2013. Antimicrobial activity of metals: mechanisms, molecular targets and applications. Nat Rev Microbiol 11:371– 384.

26. Seiler C, Berendonk TU. 2012. Heavy metal driven co-selection of antibiotic resistance in soil and water bodies impacted by agriculture and aquaculture. Front Microbiol 3:399.

27. Deng S, Cai C, Wang J, Qin D, Yu L, Wang J, Dai S, Fan J, Zhang C, Li L, Song W, Hou X. 2025. Genomics and biodegradation properties of an oleophilic bacterium isolated from shale oil sludge. International Biodeterioration & Biodegradation 200:106028.

28. Puangseree J, Jeamsripong S, Prathan R, Pungpian C, Chuanchuen R. 2021. Resistance to widely-used disinfectants and heavy metals and cross resistance to antibiotics in *Escherichia coli* isolated from pigs, pork and pig carcass. Food Control 124:107892.

29. Silva I, Tacão M, Henriques I. 2021. Selection of antibiotic resistance by metals in a riverine bacterial community. Chemosphere 263:127936.

30. Li L-G. Co-occurrence of antibiotic and metal resistance genes revealed in complete genome collection. The ISME Journal.

31. Tripathi A, Jaiswal A, Kumar D, Pandit R, Blake D, Tomley F, Joshi M, Joshi CG, Dubey SK. 2025. Whole genome sequencing revealed high occurrence of antimicrobial resistance genes in bacteria isolated from poultry manure. International Journal of Antimicrobial Agents 65:107452.

32. Zhang Y-Y, Liang Z-X, Li C-S, Chang Y, Ma X-Q, Yu L, Chen L-A. 2018. Whole-Genome Analysis of an Extensively Drug-Resistant Acinetobacter baumannii Strain XDR-BJ83: Insights into the Mechanisms of Resistance of an ST368 Strain from a Tertiary Care Hospital in China. Microb Drug Resist 24:1259– 1270.

33. Ma Y, Zhang Y, Chen K, Zhang L, Zhang Y, Wang X, Xia X. 2021. The role of PhoP/PhoQ two component system in regulating stress adaptation in *Cronobacter sakazakii*. Food Microbiology 100:103851.

34. Tomy A, Yasarla R. 2025. Study of *N*-acyl homoserine lactone (AHL) degradation potential of bacteria isolated from environmental samples and their impact on quorum sensing regulated biofilm formation of *Pseudomonas aeruginosa*. Journal of Environmental Chemical Engineering 13:115974.

35. Pan Y, Zhao M, Liu W, Jia W, Li G. 2024. Study on molecular epidemiology of carbapenem resistant *Pseudomonas aeruginosa* and related genes of quorum sensing signal system. Microbial Pathogenesis 196:106899.

36. Kaesbohrer A, Bakran-Lebl K, Irrgang A, Fischer J, Kämpf P, Schiffmann A, Werckenthin C, Busch M, Kreienbrock L, Hille K. 2019. Diversity in prevalence and characteristics of ESBL/pAmpC producing *E. coli* in food in Germany. Veterinary Microbiology 233:52–60.

37. Rajput V, Minkina T, Fedorenko A, Sushkova S, Mandzhieva S, Lysenko V, Duplii N, Fedorenko G, Dvadnenko K, Ghazaryan K. 2018. Toxicity of copper oxide nanoparticles on spring barley (*Hordeum sativum* distichum). Science of The Total Environment 645:1103–1113.

38. 2023. Versatile allelic replacement and self-excising integrative vectors for plasmid genome mutation and complementation. Microbiology Spectrum 12.

39. Nolivos S, Cayron J, Dedieu A, Page A, Delolme F, Lesterlin C. 2019. Role of AcrAB-TolC multidrug efflux pump in drug-resistance acquisition by plasmid transfer. Science 364:778–782.

40. Lloyd NA, Nazaret S, Barkay T. 2018. Whole genome sequences to assess the link between antibiotic and metal resistance in three coastal marine bacteria isolated from the mummichog gastrointestinal tract. Marine Pollution Bulletin 135:514–520.

## References

1. Argudín MA, Butaye P. 2016. Dissemination of metal resistance genes among animal methicillin-resistant coagulase-negative Staphylococci. Research in Veterinary Science 105:192–194.

2. Cavaco LM, Hasman H, Aarestrup FM, Members Of Mrsa-Cg:, Wagenaar JA, Graveland H, Veldman K, Mevius D, Fetsch A, Tenhagen B-A, Concepcion Porrero M, Dominguez L, Granier SA, Jouy E, Butaye P, Kaszanyitzky E, Dán A, Zmudzki J, Battisti A, Franco A, Schwarz S, Gutierrez M, Weese JS, Cui S, Pomba C. 2011. Zinc resistance of Staphylococcus aureus of animal origin is strongly associated with methicillin resistance. Veterinary Microbiology 150:344–348.

3. Trajanovska S, Britz ML, Bhave M. Detection of heavy metal ion resistance genes in Gram-positive and Gram-negative bacteria isolated from a lead-contaminated site.

4. 2014. ISSN 2320-5407 International Journal of Advanced Research (2014), Volume 2, Issue 3, 227–233. International Journal of Advanced Research 2.

5. Rensing C, Mitra B, Rosen BP. 1997. The *zntA* gene of *Escherichia coli* encodes a Zn(II)-translocating P-type ATPase. Proc Natl Acad Sci USA 94:14326–14331.

6. Denayer S, Matthijs S, Cornelis P. 2007. Pyocin S2 (Sa) Kills *Pseudomonas aeruginosa* Strains via the FpvA Type I Ferripyoverdine Receptor. J Bacteriol 189:7663–7668.

7. Yang S, Deng W, Liu S, Yu X, Mustafa GR, Chen S, He L, Ao X, Yang Y, Zhou K, Li B, Han X, Xu X, Zou L. 2020. Presence of heavy metal resistance genes in Escherichia coli and Salmonella isolates and analysis of resistance gene structure in E. coli E308. Journal of Global Antimicrobial Resistance 21:420–426.

8. Kamika I, Momba MN. 2013. Assessing the resistance and bioremediation ability of selected bacterial and protozoan species to heavy metals in metal-rich industrial wastewater. BMC Microbiol 13:28.

9. Fierros-Romero G, Gómez-Ramírez M, Arenas-Isaac GE, Pless RC, Rojas-Avelizapa NG. 2016. Identification of *Bacillus megaterium* and *Microbacterium liquefaciens* genes involved in metal resistance and metal removal. Can J Microbiol 62:505–513.

10. Abou-Shanab RAI, Van Berkum P, Angle JS. 2007. Heavy metal resistance and genotypic analysis of metal resistance genes in gram-positive and gram-negative bacteria present in Ni-rich serpentine soil and in the rhizosphere of Alyssum murale. Chemosphere 68:360–367.

11. Borremans B, Hobman JL, Provoost A, Brown NL, Van Der Lelie D. 2001. Cloning and Functional Analysis of the *pbr* Lead Resistance Determinant of *Ralstonia metallidurans* CH34. J Bacteriol 183:5651–5658.

